# Translational isoforms of mammalian heterochromatin protein 1 (HP1) tune its interactions with chromatin effectors

**DOI:** 10.64898/2026.10.13.758779

**Authors:** Eric M. Smith, Ekaterina Khalizeva, Sofia Haug, Iain Cheeseman

## Abstract

The heterochromatin protein 1 (HP1) protein family maintains heterochromatin by reading H3K9 methylation and recruiting diverse chromatin effectors. Multiple effectors compete for the same binding surface on HP1, requiring cells to regulate HP1’s ability to engage the right effectors across differing conditions. Here, we show that mammalian cells produce multiple N-terminally extended isoforms of HP1β from conserved, in-frame near-cognate start codons in the *CBX1* 5’ UTR. Each of these isoforms retains the localization to H3K9me3 marks. However, the extended HP1β isoforms shift its binding preference away from proteins with a conserved PxVxL motif and toward the G9a/GLP complex. Structural predictions suggest that the extension blocks the PxVxL interaction surface. Indeed, we find that an extended HP1β isoform fails to bind a model PxVxL-containing substrate *in vitro*. Together, our data reveal an alternative HP1β isoform that reshapes its effector binding preferences, providing an additional regulatory layer for heterochromatin organization.

## Introduction

As cells divide, differentiate, and respond to their environment, they must tune their gene expression programs, including through large-scale changes in genome organization by adjusting chromatin states. This dynamic process requires the machinery that reads and remodels chromatin to be controlled temporally, spatially, and across physiological conditions. The heterochromatin protein 1 (HP1) family of proteins is a key player in genome organization. This conserved protein family functions to “read” specific chromatin marks and recruit downstream effectors that act to establish, spread, and maintain heterochromatin^1^. To achieve these activities, HP1 proteins have two conserved domains: an N-terminal chromodomain (CD) that recognizes and binds to di- and tri-methylated histone H3 lysine 9 (H3K9me2/3) and a C-terminal chromoshadow domain (CSD) that is responsible for homodimerization and forms a binding pocket to interact with specific chromatin effectors^1-8^. These two structured domains are connected by a disordered linker termed the hinge region and are flanked by N- and C-terminal extensions (NTE and CTE). This architecture allows HP1 proteins to simultaneously bridge nucleosomes and recruit a diverse set of chromatin effectors to heterochromatin, including histone methyltransferases, transcription factors, centromeric proteins, and both the transcription and replication machinery ^9-15^. Many of these effector proteins interact with HP1 by binding directly to the hydrophobic pocket created by the CSD-CSD dimerization interface using a short, hydrophobic PxVxL motif^5,16^. Importantly, because these effector molecules are involved in a diverse set of processes, cells must ensure these effectors interact with HP1 proteins at the correct time and in the right places within the genome.

Cells use several strategies to modulate HP1 function. First, mammals express three separate HP1 paralogs, *CBX1* (HP1β), *CBX3* (HP1γ), and *CBX5* (HP1α). Despite having identical domain architecture and a high sequence similarity between their CD and CSD, these HP1 proteins have distinct cellular roles ^8,17^. These non-overlapping functions are illustrated by their differential requirements in mice, with the loss of HP1β causing perinatal lethality whereas hypomorphic HP1γ alleles result in spermatogenesis defects ^18,19^. Second, post-translational modifications tune HP1 function to regulate where and when HP1 localizes to different genomic regions across different cell states and conditions. For example, the SUMOylation of the HP1α hinge region directs the protein to pericentromeric heterochromatin, whereas phosphorylation of HP1β at T51 releases the protein from heterochromatin in response to DNA damage ^20,21^.

Here, we assess a previously unexplored layer of HP1 protein regulation through the production of alternative protein isoforms. Alternative translation provides a mechanism to diversify the proteome by generating multiple, related protein products from a single mRNA^22,23^. Initiation at non-canonical start sites that are in-frame with the annotated start site for a protein can produce translational isoforms with distinct biochemical and cellular properties. This includes protein variants with unique signal sequences that alter cellular localization, stabilizing or destabilizing sequences that alter a protein’s half-life, or inclusion or exclusion of a domain that alters interaction partners^24-26^.

Our results reveal a new mechanism by which cells control which chromatin effectors interact with HP1β. We find that human *CBX1* encodes multiple N-terminally extended HP1β isoforms that are translated from in-frame non-AUG start codons in the 5’ untranslated region (UTR). Each of these isoforms can be recruited to H3K9me3 marks in cells. However, the extended isoforms alter the preference of HP1β for its downstream effectors, redirecting HP1β away from PxVxL motif-dependent effectors, such as the CAF-1 complex, and toward the G9a/GLP complex. Structural predictions suggest that the extension residues interact with the hydrophobic pocket created by the CSD-CSD dimerization interface, creating a steric block for chromatin effectors that use a PxVxL motif to bind to HP1β. Consistent with this model, the extended HP1β isoform no longer binds to a PxVxL-containing model peptide *in vitro*. Together, this work reveals that cells use alternative translation initiation to produce HP1 isoforms with distinct effector preferences, providing cells with a powerful regulatory layer to tune HP1 function as cellular demands change.

## Results

### Identification of conserved N-terminally extended HP1β isoforms

Although each mRNA is typically considered to generate a single protein product, we sought to identify protein variants generated by alternative translation initiation that have the potential to impact genome organization and function. In analyzing start site ribosome profiling data from diverse sources, we identified N-terminally extended isoforms of heterochromatin protein 1 (HP1) that use upstream, in-frame non-canonical start sites. In particular, start-site ribosome profiling identified in-frame 4, 41, and 54 amino acid extensions of human HP1β (*CBX1*) (Figure 1A and 1B)^27^. Interestingly, analysis of start site ribosome profiling data from mouse embryonic stem cells also revealed the presence of 4 and 42 amino acid N-terminal extensions for mouse HP1β (*CBX1*) (Figure 1C and 1D)^23^.

**Figure 1.**
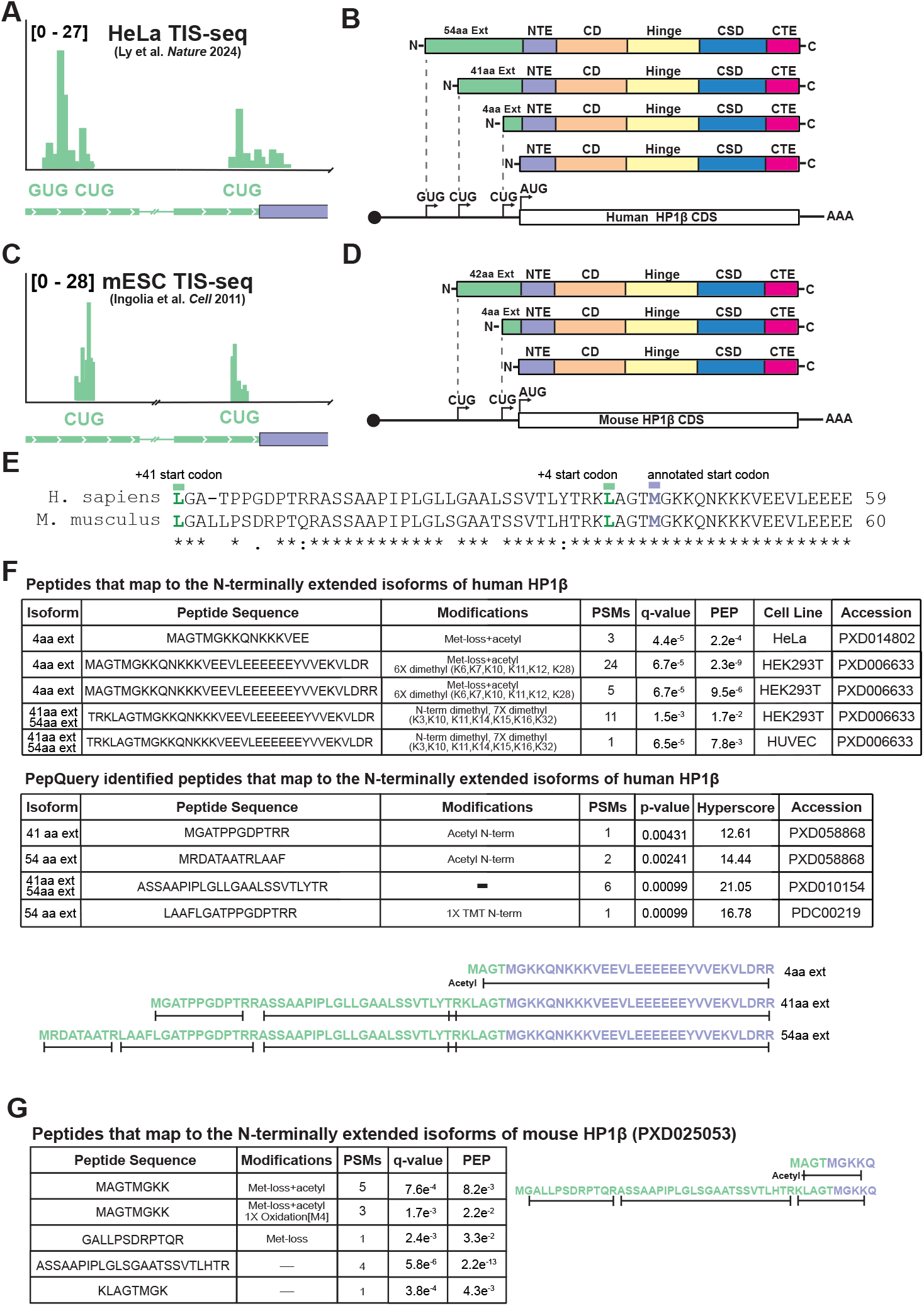
Multiple HP1β isoforms can be produced from human and mouse *CBX1* mRNA. **(A)** Translation initiation site ribosome profiling read distribution at human *CBX1* alternative start codons. The green peaks are reads at the alternative start sites in the 5’ UTR of the transcript. **(B)** A cartoon schematic depicting the proteins produced from alternative translation initiation along with domain diagrams for each isoform. **(C)** Translation initiation site ribosome profiling read distribution at mouse *CBX1* alternative start codons. The green peaks are reads at the alternative start sites in the 5’ UTR of the transcript. **(D)** A cartoon schematic depicting the proteins produced from alternative translation initiation along with domain diagrams for each isoform. **(E)** A sequence alignment created using Clustal Omega between the 41 residue N-terminal extension of human and the 42 residue extension of mouse HP1β. The residues marked in green are the sites of alternative translation initiation. The amino acid identity of the marked non-canonical start codons matches the coding sequence for the gene, but would be substituted for methionine when used as a translation initiation site. **(F)** Tables showing peptides mapping to the indicated HP1β isoform. Peptides in the top table were identified by reprocessing data in Proteome Discoverer. Peptides in the bottom table were identified by search with PepQuery^41^. The accession number refers to the PRoteomics IDEntifications database (PRIDE) or Proteomic Data Commons (PDC) number. The sequences below highlight where in the sequence identified peptides map. **(G)** A table showing peptides mapping to the indicated HP1β isoform. Peptides were identified by reprocessing data in Proteome Discoverer. All the mouse peptides came from PXD025053^30^.

Unlike coding regions that are under tight selection for their protein coding potential, non-coding 5’ UTR regions typically diverge substantially between species. In contrast, consistent with the presence of protein-coding alternate extensions, we found that the 5’ UTR region upstream of the annotated CBX1 start codon is highly conserved across mammals and maintains its protein coding potential. Indeed, the 41 and 42 amino acid extension sequences from humans and mice, respectively, display 80% amino acid identity (Figure 1E). We also detected a strong conservation of these N-terminal extensions, including the presence of near-cognate start codons amongst diverse mammalian species (Figure S1A). Finally, in addition to CBX1, based on start site ribosome profiling data, we identified conserved translational extensions for human HP1α (*CBX5*) suggesting that this may be a property of diverse HP1 isoforms (Figure S1B).

To obtain orthologous evidence for the production of these alternative translational isoforms, we analyzed publicly available mass spectrometry data sets ^28-31^. We identified peptides corresponding to each HP1β translational isoform in human cells, including N-terminal peptides for each isoform (Figure 1F and S1C-E). Similarly, we identified peptides uniquely corresponding to both the 4 and 42 amino acid extensions of mouse CBX1 (Figure 1G and S1F-H). Together, these data indicate that cells produce evolutionarily-conserved alternate translational isoforms for HP1β (*CBX1*) with the potential to impact its roles in heterochromatin organization.

### N-terminally extended HP1β isoforms display modified chromatin localization

Alternative translational isoforms of annotated proteins can modulate protein properties to impart distinct cellular behaviors from their annotated counterparts, including changes to their subcellular localization^24^. To test if the N-terminally extended CBX1 isoforms alter HP1β localization, we compared the localization of each HP1β isoform to the annotated HP1β protein. For these experiments, we generated human HeLa or mouse NIH3T3 cell lines that ectopically express a C-terminally GFP-tagged version of each protein. Consistent with the established localization of HP1β, we found that the annotated HP1β protein and each alternative translational isoform localized to the nucleus. In each case, we observed a diffuse nuclear signal as well as the presence of nuclear foci (Figure 2A and S2A).

**Figure 2.**
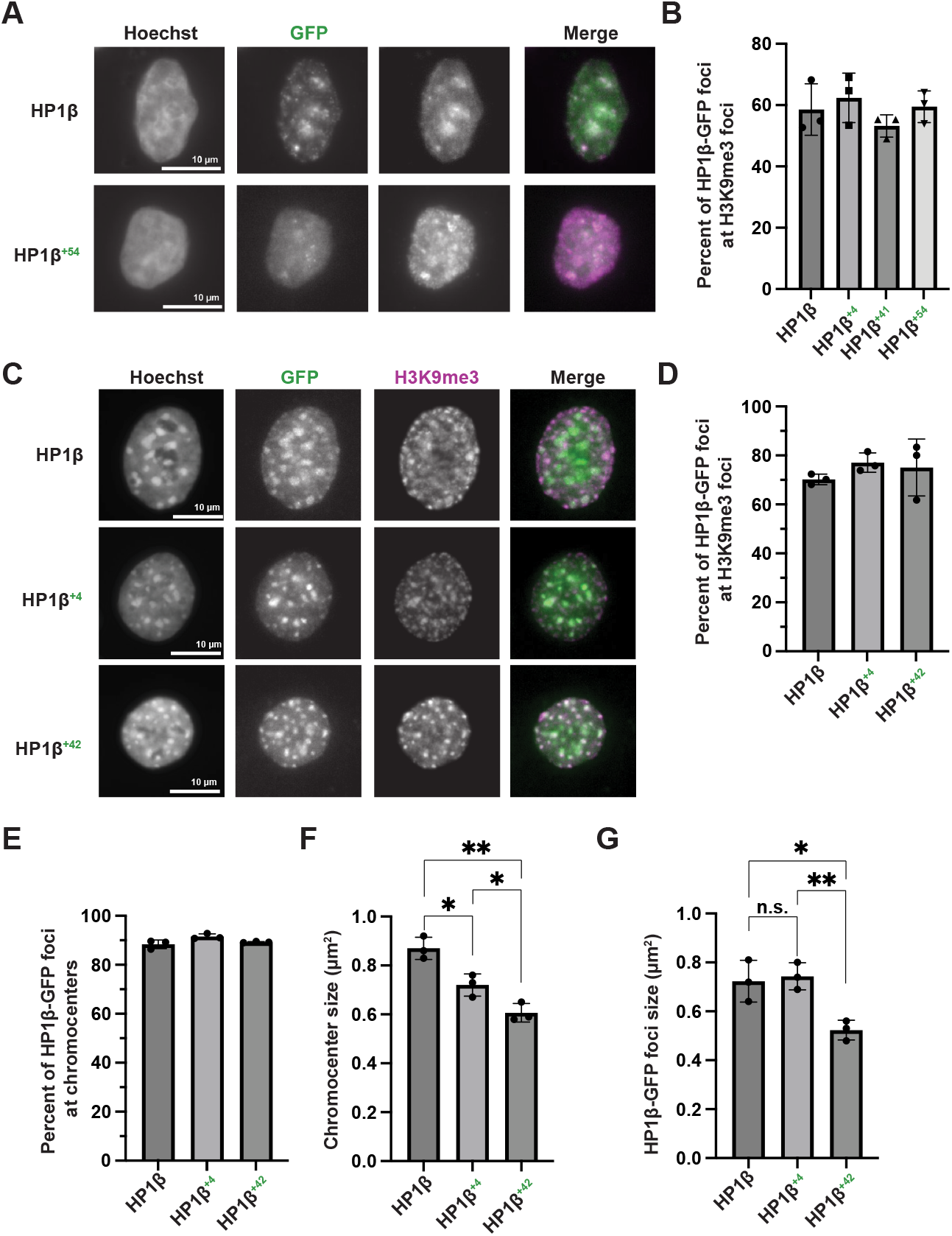
Isoforms of HP1β have differential chromatin localization. **(A)** Representative images of a single 2 µm equatorial slice of HeLa cells ectopically expressing the indicated C-terminally GFP tagged HP1β construct. The Hoechst and H3K9me3 channels are scaled identically, but the GFP channels are scaled individually. **(B)** A bar graph showing the percentage of HP1β-GFP foci that overlap with H3K9me3 foci in HeLa cells. The plotted value is the mean and the error bars represent the standard deviation from three biological replicates. **(C)** Representative images of a single 2 µm equatorial slice of NIH3T3 cells ectopically expressing the indicated C-terminally GFP tagged HP1β construct. The Hoechst and H3K9me3 channels are scaled identically, but the GFP channels are scaled individually. **(D)** A bar graph showing the percentage of HP1β-GFP foci that overlap with H3K9me3 foci in NIH3T3 cells. The plotted value is the mean and the error bars represent the standard deviation from three biological replicates. **(E)** A bar graph showing the percentage of HP1β-GFP foci that overlap with chromocenters in NIH3T3 cells. The plotted value is the mean and the error bars represent the standard deviation from three biological replicates. **(F)** A bar graph showing the size (µm^2^) of chromocenters in NIH3T3 cells. The plotted value is the mean and the error bars represent the standard deviation from three biological replicates. p-values were calculated using a two-tailed Welch’s t-test. The p-value for the comparison of HP1β^+42^ to HP1β is 0.0018, HP1β^+42^ to HP1β^+4^ is 0.035, and HP1β^+4^ to HP1β is 0.016. *p < 0.05, and **p < 0.005. **(G)** A bar graph showing the size (µm^2^) of HP1β-GFP foci in NIH3T3 cells. The plotted value is the mean and the error bars represent the standard deviation from three biological replicates. p-values were calculated using a two-tailed Welch’s t-test. The p-value for the comparison of HP1β^+42^ to HP1β is 0.0377, HP1β^+42^ to HP1β^+4^ is 0.0065, and HP1β^+4^ to HP1β is 0.752. *p < 0.05, and **p < 0.005.

Because HP1β contains a chromodomain that is responsible for binding to trimethylated histone H3K9 (H3K9me3; ^2^), we next tested if these puncta co-localize with H3K9me3 heterochromatin. For the annotated HP1β protein, we found that 59% of HP1β-GFP co-localized with H3K9me3 marks in HeLa cells. Similarly, 62% of HP1β^+4^, 53% of HP1β^+41^, and 60% of HP1β^+54^ co-localized with H3K9me3 marks (Figure 2B and S2B). Similarly, for the mouse HP1β isoforms, we found that HP1β^+42^, HP1β^+4^, and HP1β foci overlapped with H3K9me3 marks (with 75%, 77%, and 70% co-localization, respectively; Figure 2C-D and S2C). Thus, the presence of the translational extension does not disrupt the recruitment of the HP1 chromodomain to H3K9me-containing chromatin.

Mouse cells contain heterochromatin-dense structures, termed chromocenters, which are marked by bright Hoechst foci in the nucleus (Figure 2C;^32^). We found that both the annotated and extended mouse HP1β isoforms co-localized with chromocenters, with 88-92% of puncta formed by each isoform of HP1β overlapping with a chromocenter (Figure 2E and S2D). Interestingly, when we measured the size of the chromocenters, we found that cells ectopically expressing annotated HP1β had the largest chromocenters (0.87 ± 0.05 µm^2^), whereas cells ectopically expressing HP1β^+4^ (0.72 ± 0.05 µm^2^), or HP1β^+42^ (0.61 ± 0.04 µm^2^) had significantly smaller chromocenters (Figure 2F and S2E). The size of the GFP puncta showed a similar trend, with annotated HP1β (0.72 ± 0.09 µm^2^) and HP1β^+4^ (0.74 ± 0.06 µm^2^) forming larger puncta than HP1β^+42^ (0.52 ± 0.05 µm^2^) (Figure 2G and S2F). Thus, although each HP1β isoform is recruited to chromocenters, cells that express HP1β^+42^ form smaller chromocenters suggesting that this protein may act to impact chromocenter formation beyond H3K9me3 binding. As chromocenter formation requires the action of chromatin effectors, such as CAF-1, that establish heterochromatin, along with higher-order interactions that cluster repetitive DNA elements, this size difference may reflect reduced heterochromatin spreading or altered clustering of pericentromeric DNA^33-35^. Together, these results suggest that the recruitment of the HP1β chromodomains to H3K9me3 chromatin is retained in the alternative isoforms, but that the extended HP1β isoforms may have altered properties.

### N-terminally extended HP1β isoforms display differential protein interactions

We next considered whether the N-terminal translational HP1β isoforms alter protein interaction interfaces or its binding partners. HP1β is an obligate homodimer such that the ectopically expressed GFP fusion constructs have the potential to dimerize with the endogenous HP1 proteins. Thus, to test the interaction partners for each isoform individually, we first created HeLa cell lines in which endogenous *CBX1* was eliminated using CRISPR-Cas9 gene targeting, followed by ectopic expression of GFP fusions for either annotated HP1β or HP1β^+54^ (Figure S3A and B). Based on quantitative immunoprecipitation-mass spectrometry (IP-MS) experiments relative to a GFP control IP, we found that both HP1β and HP1β^+54^ interacted with established chromatin effectors, including the CAF-1 complex (CHAF1A, CHAF1B, and RBBP4), the G9a/GLP complex (EHMT1, EHMT2, WIZ, and ZNF644), CHAMP1, NIPBL, and TRIM24 (Figure S3C, S3D, and Supplemental Table 1). Thus, the overall binding capacity of HP1 for downstream effectors remains intact in the alternative translational isoform.

Interestingly, when we compared the HP1β and HP1β^+54^ IPs, we found that HP1β^+54^ preferentially interacted with G9a/GLP components, with a 2-4 fold increase in G9a/GLP subunits. Reciprocally, we found that the annotated HP1β isoform had a preference for interactions with PxVxL containing chromatin effectors such as the CAF-1 complex, NIPBL, and TRIM24, with a 2-5 fold enrichment (Figure 3A and 3B). To test this further, we additionally performed TMT-based quantitative mass spectrometry experiments in a HeLa background containing the endogenous HP1 proteins. In this case, we also found that HP1β^+54^ isolated more of the G9a/GLP complex (EHMT1, EHMT2, ZNF644, and WIZ), but lower amounts of PxVxL containing effectors such as the CAF-1 complex (CHAF1A, CHAF1B, and RBBP4), ADNP, CHD4, POGZ, and TRIM28 than the annotated protein (Figure 3C, 3D, and S3E), although to a slightly reduced extent relative to the complete replacement conditions. Finally, we performed TMT-based quantitative mass spectrometry experiments in mouse NIH3T3 cells and similarly observed that the mouse HP1β^+42^ isoform preferentially interacted with the G9a/GLP complex components (Figure S3F-H).

**Figure 3.**
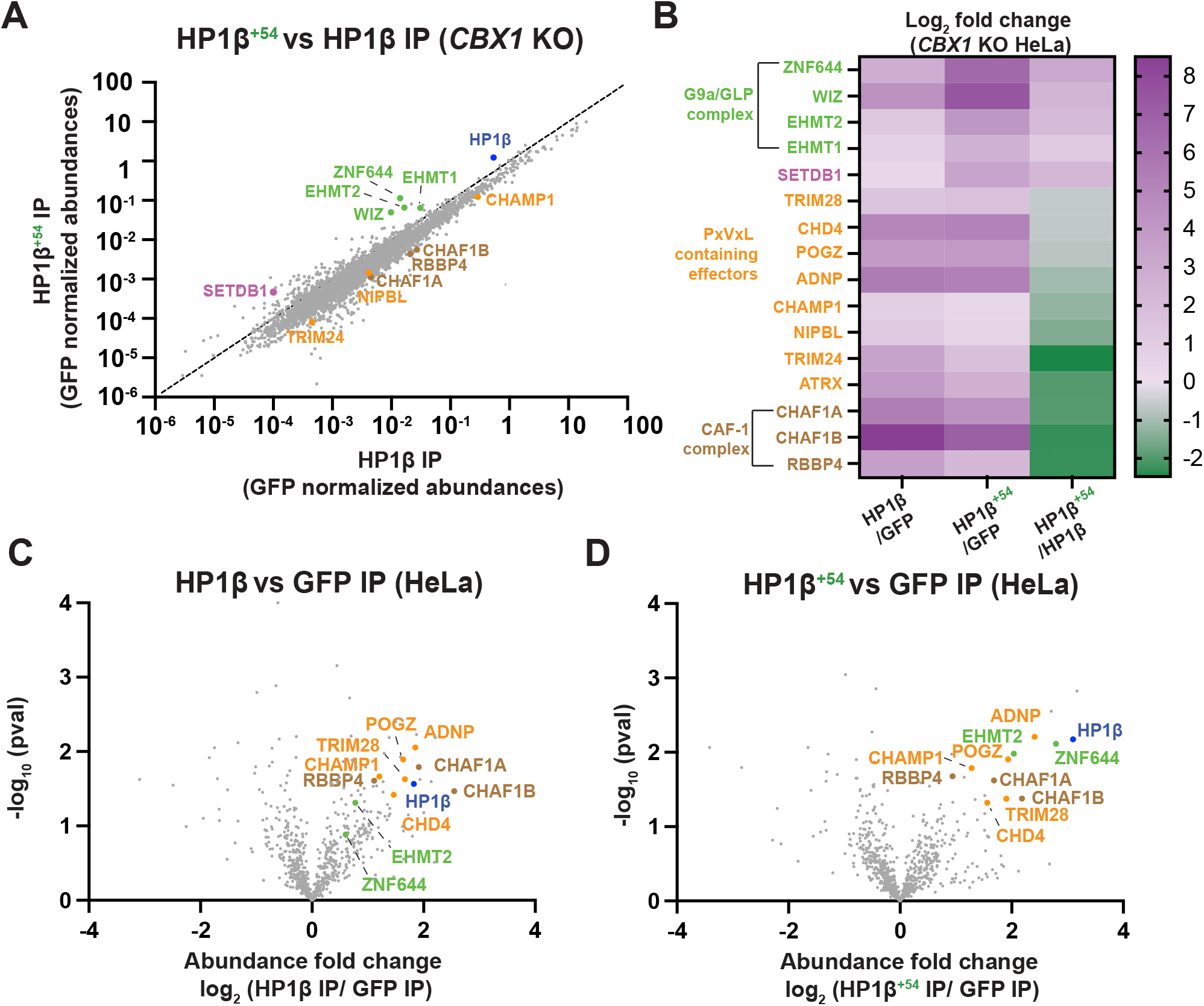
Quantitative mass spectrometry reveals preferential interactions between HP1β isoforms and chromatin effectors. **(A)** Scatter plot showing the GFP normalized abundance of the proteins detected in the HP1β^+54^ IP compared to the HP1β IP from HeLa cells in which *CBX1* was targeted using CRISPR/Cas9. HP1β is highlighted in blue, components of the G9a/GLP complex are shown in green, PxVxL containing proteins are shown in orange, and the CAF-1 complex subunits are shown in brown. **(B)** A heatmap showing the log_2_ fold abundance change of selected proteins in the indicated IP experiments. The HP1β^+54^/ HP1β values are GFP normalized. **(C)** Volcano plot showing the log_2_ fold change (median abundance normalized) of HP1β IP/GFP IP TMT-based proteomics experiments. **(D)** Volcano plot showing the log_2_ fold change (median abundance normalized) of HP1β^+54^ IP/GFP IP from TMT-based proteomics experiments.

Together, these data indicate that the extended HP1β protein has distinct preferences for its downstream effectors in cells, with increased preference for the G9a/GLP methyltransferase complex and a decreased binding to PxVxL-containing effectors.

### The N-terminal HP1β extension forms a predicted auto-inhibitory interaction

One potential explanation for the shift in binding preferences that we observed in the IP-mass spectrometry analysis is that the residues that comprise the N-terminal extension could alter the structure of HP1β or add a new interaction domain. To evaluate this, we first performed structural predictions of HP1β homodimers for each isoform using AlphaFold (Figure 4A, 4B, and S4A-F). Each HP1β monomer within the homodimer is predicted to have two folded domains; a chromodomain (CD) and a chromoshadow domain (CSD) (Figure 4A), consistent with prior structural analyses^1,5,36^. These two structured domains are connected by a long linker created by the hinge domain. Homodimers of the alternative translational isoforms were predicted to have an overall similar architecture to the dimers composed of the annotated protein, consistent with the ability of the chromodomain to be recruited to H3K9me3-containing chromatin.

**Figure 4.**
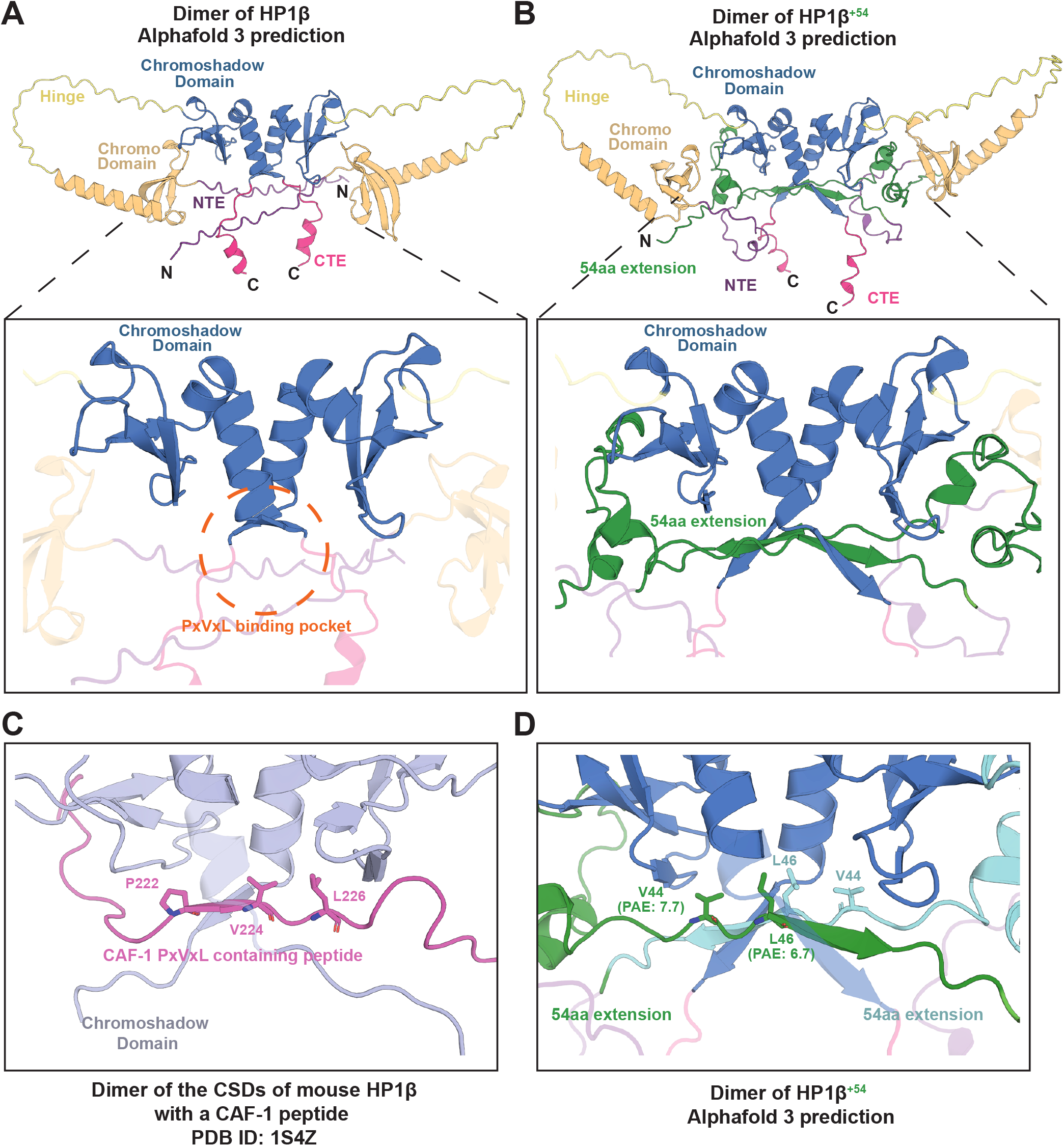
Structural predictions show an intramolecular interaction between the HP1β N-terminal extension and its effector binding groove. **(A)** A cartoon representation of the predicted structure of a dimer of annotated human HP1β that was created using AlphaFold 3^42^. Each chain of the dimer has the same coloring scheme. For each chain the N-terminal extension (NTE) is shown in purple, the chromodomain is shown in orange, the hinge region is shown in yellow, the chromoshadow domain is shown in blue, and the C-terminal extension is shown in pink. The bottom panel is a zoomed in view of the hydrophobic binding pocket that is formed by the CSD-CSD mediated homodimerization interface. **(B)** A cartoon representation of the predicted structure of the dimer of HP1β^+54^ that was created using AlphaFold 3. The color scheme is the same as (A) except the residues that are appended to the N-terminus by the translational isoform are shown in green. The bottom panel highlights the interaction the extended isoform is predicted to make with the hydrophobic pocket. **(C)** A magnified view of the PxVxL-CSD interaction from PDB ID: 1S4Z^5^. **(D)** A magnified view of (B) highlighting the VxL like motif in HP1β^+54^.

For the annotated protein, the N- and C-terminal extensions (NTE and CTE) are predicted to be largely unstructured and protrude out around the dimerization interface formed by the two chromoshadow domains, leaving the PxVxL binding pocket exposed (Figure 4A-D: bottom panel). Interestingly, we observed a distinct predicted behavior for the human HP1β^+54^, human HP1β^+41^, and mouse HP1β^+42^ isoforms (Figure 4B, S4C, and S4G). In each case, the predictions suggest that residues from the extension fold back to interact with the hydrophobic pocket created by the CSDs (Figure 4B, S4C, and S4G: bottom panel). This has the potential to significantly affect HP1 binding partners that associate with the CSD in this region, such as through PxVxL motifs. In contrast, the residues that make up the extension of HP1β^+4^ are not predicted to contact the CSD (Figure S4D).

To evaluate this predicted contact between the translational extension and the CSD, we inspected the predicted aligned error (PAE) between the extended residues and rest of the protein. Most of the extension region displayed high PAE (low confidence) relative to the folded domains (Figure S4B). However, the extension residues contacting the hydrophobic pocket showed low PAE (high confidence) with respect to the CSD pocket residues (Figure 4D). Additionally, these residues are hydrophobic (VxL), and predicted to engage the CSD binding pocket in a similar manner to the solved structure of a PxVxL containing peptide of CAF-1 (Figure 4C and 4D). Together, the structural predictions and IP-MS data suggest a model in which the longer HP1β extensions act through an auto-inhibitory mechanism to alter interactions between HP1β and downstream chromatin effectors, shifting the binding preferences of the longer HP1β isoforms.

### N-terminally extended HP1β isoforms display altered binding to a chromatin effector substrate *in vitro*

The structural predictions described above suggest that the extended HP1β could alter binding to its effector proteins through the chromoshadow domain (Figure 5A). However, the chromoshadow domain (CSD) is also required for HP1β homodimerization. To test this structural model, we reconstituted each HP1β isoform by expression in bacteria. In each case, the purified proteins behaved as a dimer, suggesting that the CSD homodimerization activity is intact (Figure S5A-G).

**Figure 5.**
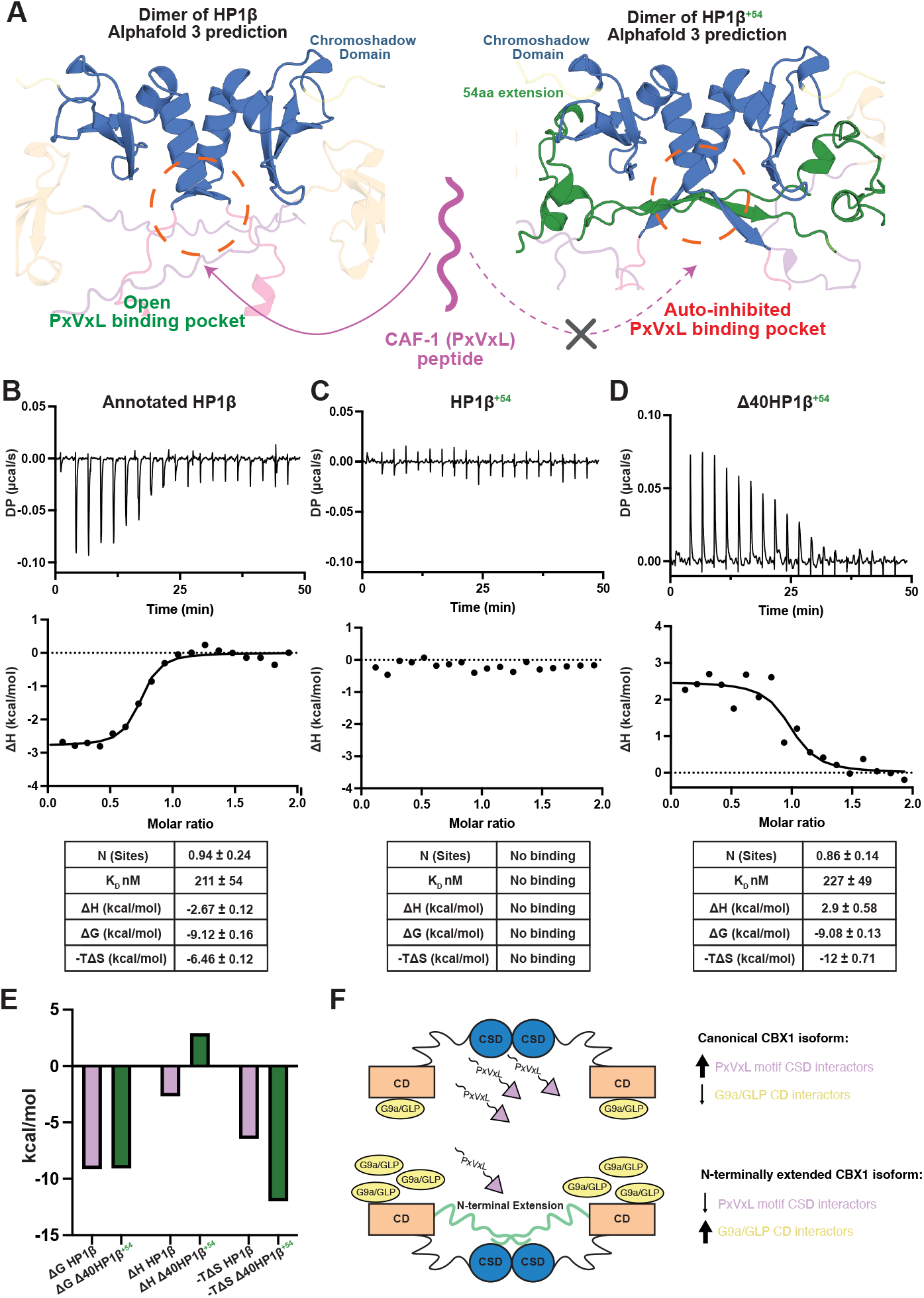
N-terminally extended isoforms of HP1β fail to bind to PxVxL containing peptide. **(A)** A schematic showing the experimental setup. **(B)-(D)** Representative raw thermograms (top) and integrated heats fit to a single-site model (bottom). For each panel a CAF-1 peptide^4^ was titrated into the cell containing the indicated HP1β construct. The table below each shows the mean and standard deviation from three replicate experiments. **(E)** A bar graph highlighting how CAF-1 binding to HP1β is enthalpically favorable, but binding is unfavorable to the Δ40HP1β^+54^ construct. **(F)** A cartoon depicting the binding preferences for each HP1 isoform.

Prior structural studies found that a hydrophobic pocket formed at the interface of the two CSDs creates a binding platform for diverse chromatin effectors, many of which contain a PxVxL motif^5,16^. For example, a PxVxL-containing peptide of the chromatin effector CAF-1 binds to the CSDs of HP1α with an affinity of 430 nM^37^. To test the binding of CAF-1 to the extended isoforms of HP1β, we performed isothermal titration calorimetry (ITC) on the purified HP1β constructs (Figure 5B-C and S5H). We found that annotated HP1β had an affinity of 211 nM for a 20 amino acid CAF-1 peptide, about two-fold higher affinity compared to the CSDs of HP1α (Figure 5B). Strikingly, we found that there was no detectable binding of the CAF-1 peptide to the HP1β^+54^ construct (Figure 5C). Thus, the presence of the extension residues disrupts binding to this well-defined PxVxL substrate, consistent with structural predictions. To test the structural model (Figure 5A), we next expressed and purified truncated HP1β^+54^ constructs and performed additional ITC experiments (Figure 5D and S5I-K). We found that truncating 10, 20, or 30 residues from the extended protein isoform did not rescue the ability of HP1β to bind to the CAF-1 peptide (Figure S5I-K). However, further truncation of 40 residues (leaving just 14 amino acids appended to the annotated isoform) rescued binding of CAF-1 to the N-terminally extended HP1β, with an affinity matching CAF-1 for the annotated HP1β protein (Figure 5D). Notably, although the change in Gibbs free energy (ΔG) was consistent between the annotated and Δ40HP1β^+54^, the binding changed from being exothermic to endothermic (Figure 5E). To offset this change, there was a major shift in ΔS, suggesting that a conformational change accompanies CAF-1 binding (Figure 5E). Together, these results suggest that the N-terminal extension has auto-inhibitory activity that alters its protein-protein interactions, reducing the binding to effectors that interact through a canonical PxVxL motif (Figure 5F).

## Discussion

Here, we identify alternative translational isoforms for the heterochromatin protein HP1β, providing a previously unrecognized strategy for HP1β regulation. We found that mammalian cells produce multiple N-terminally extended HP1β isoforms from non-canonical upstream start codons for both human and mouse *CBX1*. These isoforms are recruited to H3K9me3-containing chromatin, but display different preferences for chromatin effector binding. In particular, they display reduced binding to PxVxL-containing chromatin effectors, such as the CAF-1 complex, both in cells and using recombinant proteins, but an increased preference for the G9a/GLP complex. Structural predictions and biochemical experiments suggest that the residues that make up the extension fold back onto the hydrophobic pocket created by the dimerization of the HP1β chromoshadow domains. Together, these data create a model in which the production of alternative HP1β isoforms from the same mRNA may provide cells with a mechanism to control which chromatin effectors can interact with HP1β, providing a powerful strategy for genome organization across physiological conditions. Indeed, differential translation across conditions could result in a distinct balance of HP1 isoforms, acting to reshape aspects of genome organization^24,25,27^.

Previous work has focused on defining how HP1 function can be fine-tuned across different physiological circumstances, including through the production of paralogs and by post-translational modifications^8,38^. Alternative translation initiation is one axis that has yet to be explored for the fine-tuning of HP1 function. The differential decoding of individual mRNAs by alternative translation initiation gives cells another axis to adapt core cellular processes, producing proteins with altered biochemical and cellular properties. Here we describe another way alternative translation initiation can be used to alter protein function, by adding residues onto a protein’s N-terminus that serve as an auto-inhibitory regulator of protein-protein interactions. This finding has similarities to the auto-inhibition of the *Schizosaccharomyces pombe* HP1 protein Swi6, in which a sequence on Swi6 acts as a histone mimic, so one subunit of the Swi6 dimer blocks the chromodomain of its partner from reading methylation marks^39^. Interestingly, this regulatory interaction can be outcompeted by the simultaneous recognition of nucleosome DNA and H3K9 methylation marks^39^. Although the auto-inhibition described here is to the chromoshadow domain rather than the chromodomain, it may serve an analogous role in ensuring that HP1β interacts with the correct partners at the correct time and location in the nucleus. Our biochemical data suggest that this auto-inhibition is difficult to relieve with just one peptide substrate. However, our IP-MS experiments show that HP1β^+54^ interacts with several PxVxL containing proteins, suggesting that this auto-inhibition may be relieved in some circumstances or that these interactions may be retained indirectly by shared chromatin interactions. If auto-inhibition is relieved, it may be through a similar mechanism to Swi6, or may be regulated by an alternative mechanism such as a post-translational modification.

Either mechanism would make this a tunable switch that could provide a powerful way for cells to fine-tune which proteins HP1β interacts with across different physiological conditions. Although we focused on HP1β here, the detection of translational isoforms of other chromatin regulators, such as HP1α, suggest that alternative translation initiation may be a more prevalent mechanism to regulate chromatin than previously appreciated.

## Acknowledgments

The authors thank the members of the Cheeseman laboratory for thoughtful discussion and assistance. Specifically, we want to thank Jimmy Ly for critical work that enabled this study and for thoughtful discussions throughout this project. The authors would like to thank the MIT Biophysical Instrumentation Facility (BIF) for ITC, the Whitehead Flow Cytometry Core for cell sorting and analysis, and the Whitehead Quantitative Proteomics Core for help and support with mass spectrometry. This work was supported by grants from NIGMS (R35GM126930), the Chan Zuckerberg Initiative (Rare as One), and MIT (Bose Fellowship) to IMC, an MIT HEALS graduate fellowship to EK, and a post-doctoral fellowship from the American Cancer Society to EMS.

## Experimental Procedures

### Plasmid Constructs

To clone all constructs used in this study the sequence of human and mouse *CBX1* was synthesized by TWIST with one mutation, a silent mutation was inserted to remove the XhoI site from the construct. The human 54aa extension of HP1β, the mouse 42aa extension of HP1β, the human 41 aa extension of HP1β, human/mouse 4aa extension, and human/mouse annotated protein were amplified from these oligomers and restriction cloning was used to insert each construct into a pBABEblast-GFP fusion protein vector. Restriction cloning was used to insert each construct into pGEX-6P-1 for expression in *E. coli*. All single-guide RNAs (sgRNAs) were cloned into pX330 (Addgene 42230). The following sgRNA sequence was used to target human *CBX1*: 5’-GCAGAAACTCAGCAATGAGGT-3’.

### Tissue culture

HeLa, HEK293-GP, and NIH3T3 cells were cultured in Dulbecco’s modified Eagle medium (DMEM) supplemented with 10% tetracycline-free fetal bovine serum, 100 U/ml penicillin–streptomycin and 2 mM L-glutamine at 37 °C with 5% CO_2_.

### Cell-line generation

Cell lines expressing C-terminally GFP-tagged human canonical HP1β, human 54aa extension of HP1β, human 41 aa extension of HP1β, human 4aa extension, mouse canonical HP1β, mouse 42aa extension of HP1β, mouse 4aa extension of HP1β, and GFP control cells were generated by retroviral transduction. Each retrovirus was created by transfecting a pBABEblast vector and VSV-G into HEK293-GP cells. Retroviral transduction was performed by mixing each virus with 10 µg/ml polybrene and spinning cells for 45 minutes at 2250 rpm. After 48 hours retrovirus was removed from the cells and infected cells were selected with 2 µg/ml blasticidin (Life Technologies) for 2 weeks, changing media every 4 days.

### Immunofluorescence

The indicated HeLa or NIH3T3 cell lines were seeded on poly-L-lysine coated coverslips. Pre-extraction in 0.5% Triton X-100 in 1X PBS was performed for 10 minutes before samples were fixed in 1X PBS with 4% formaldehyde and 0.1% Triton X-100 at room temperature for 10 min. Following fixation, cells were washed 3 times with 0.1% Triton X-100 in 1 X PBS (PBSTx) and blocked for 1 hour in Abdil (3% BSA, 0.1% Triton X-100, and 0.2% sodium azide in 1X TBS). Cells were incubated with anti-H3K9me3 antibody (1:1000, Active Motif, 39065) at 4°C in a humidified chamber overnight. Cells were washed with PBSTx 3 times, then incubated with GFP booster (ChromoTek, gba488), secondary antibody (1:300, Donkey Anti-Rabbit IgG (H+L) Cy3) diluted in AbDil for 1 h at room temperature. Cells were then washed 2 times with PBSTx, and 1 time with PBSTx with Hoechst. Cells were then mounted to glass cover slips with ProLong Gold antifade reagent (Invitrogen, P36390). A DeltaVision Ultra High-Resolution microscope with a 60x/1.42NA objective (Cytiva) was used to image the cells. Images were analyzed using ImageJ and Cell Profiler.

### Western blot analysis

Western blots were performed on isolated HeLa clones that had been transfected with 750 ng of pX330 containing a sgRNA to *CBX1* using X-tremeGENE 9 (Roche). Single clones were grown up and plated into a 6-well dish. Samples were lysed by the addition of 1X Laemmli sample buffer (100 mM Tris pH 6.8, 12.5% glycerol, 1% SDS, 0.1% bromophenol blue, and freshly added 200 mM β-mercaptoethanol). Samples were heated for 10 minutes at 95 °C before being separated on a 15% SDS-PAGE gel. A transblot turbo (Bio-Rad) was used to transfer the separated proteins onto a nitrocellulose membrane. The membrane was blocked for 1 h in 3% milk dissolved in 1X TBST. For immunoblotting an anti-CBX1/ HP1β antibody (Abcam: ab10478) was diluted 1:5000 and an anti-alpha tubulin (DM1A, Millipore-Sigma T9026) was diluted 1:2000 in 3% milk and incubated with the blot at 4 °C overnight. The blot was washed three times with 1X TBST for five minutes each wash before being incubated with IRDye 680RD goat anti-mouse (LI-COR 92668070) and IRDye 800CW goat anti-rabbit (LI-COR 92632211) diluted 1:10,000 in 3% milk for one hour. The blot was then washed three more times for 5 minutes in 1X TBST before being imaged on an Odyssey CLx (LI-COR).

### Immunoprecipitation-mass spectrometry

For each IP-MS experiment ten 15 cm dishes of HeLa or NIH3T3 cells ectopically expressing GFP or GFP-tagged HP1β constructs were grown to approximately 90% confluency. Cells were harvested by incubation in 5 mM EDTA in 1X PBS for 5 minutes at 37 °C. Dislodged cells were pelleted by centrifugation at 200 x g. Cells were washed once with 1X PBS, pelleted again, and then washed with 1x lysis buffer without detergent (25 mM HEPES pH 8.0, 2 mM MgCl2, 0.1 mM EDTA pH 8.0, 0.5 mM EGTA pH 8.0, 300 mM KCl and 10% glycerol). Finally, after pelleting the cells and removing the supernatant the cells were resuspended in 1x lysis buffer without detergent at a 1:1 ratio and flash frozen in liquid nitrogen before being stored at -80 °C. An equal volume of 1.5X lysis buffer detergent (37.5 mM HEPES pH 8.0, 3 mM MgCl2, 0.15 mM EDTA pH 8.0, 0.75 mM EGTA pH 8.0, 450 mM KCl, 15% glycerol and 0.225% Nonidet P-40 substitute) supplemented with 1 cOmplete EDTA-free protease inhibitor cocktail, 1 mM PMSF, 0.4 mM sodium orthovanadate, 5 mM sodium fluoride and 20 mM β-glycerophosphate was added to each frozen cell pellet before thawing the cells in a 37 °C water bath. Cells were lysed by 10 cycles of sonication at high amplitude with a 30 second on time and 1 minute off time (Bioruptor, Diagenode). The lysate was then clarified by centrifugation at 21,000 x g for 30 minutes at 4 °C. Following this spin, the supernatant was incubated with Protein A beads (Bio-Rad) coupled to a rabbit anti-GFP antibody (83 µg/ 150 µl packed beads)^40^ at 4 °C for 1 hour on a rotating wheel. The beads were then rinsed 3 times with 1X lysis buffer (25 mM HEPES pH 8.0, 2 mM MgCl2, 0.1 mM EDTA pH 8.0, 0.5 mM EGTA pH 8.0, 300 mM KCl, 10% glycerol, and 0.1% Nonidet P-40 substitute) supplemented with 1 mM DTT, 10 µg/ml leupeptin/pepstatin/chymostatin, and 0.02 U/μl of SUPERaseIn. The beads were then washed 2 times for 5 minutes on a spinning wheel in 1X lysis buffer followed by one wash in 1X lysis buffer without detergent. Next, three elutions were performed by incubating the beads with 2 bead volumes of 100 mM glycine pH 2.6 for 5 minutes at 4 °C on a rotating wheel. The three elutions were pooled and Tris pH 8.5 was added to a final concentration of 200 mM. Next, trichloroacetic acid was added to 1/5^th^ the final volume, samples were gently mixed, and the proteins were precipitated overnight at 4 °C. The precipitated proteins were pelleted by centrifugation at 21,000 x g for 30 minutes. The protein pellets were then washed three times with 0.5 ml cold acetone. Proteins were pelleted by centrifugation at 21,000 x g for 10 minutes between washes. Proteins were then dried in a SpeedVac vacuum concentrator for 5 minutes. The precipitated proteins were resuspended in 5% SDS, 50 mM triethylammonium bicarbonate pH 8.5 (TEAB), 20 mM DTT. The resuspended proteins were incubated at 95 °C for 10 minutes and then cooled to room temperature. Freshly prepared 40 mM iodoacetamide was added to the resuspended proteins and the alkylation reaction was carried out for 30 minutes in the dark. The reaction was quenched by adding 2.5% v/v phosphoric acid. Next, S-Trap binding buffer (90% MeOH, 100 mM TEAB, pH 7.55) was added to the quenched alkylation reaction and each sample was loaded onto an S-Trap microcolumn (Protifi) by centrifugation at 4000 x g for 1 minute. The eluate was passed through the column a second time and subsequently the column was washed 4 times with 150 µl of S-Trap binding buffer. An on-column digestion was performed by adding 1 µg of trypsin diluted in 50 mM TEAB pH 8.5 to the top of the column and placing the column in a humidified chamber at 37 °C overnight. The tryptic peptides were eluted in three steps: first, with 40 µl of 50 mM TEAB, followed by 40 µl of 0.2% formic acid, and finally with 35 µl 50% acetonitrile/0.2% formic acid. The eluted peptides were pooled, flash frozen, and then lyophilized. The amount of peptide obtained was determined using the Pierce fluorometric peptide assay kit (Pierce) following the manufacturer’s protocol. The lyophilized peptides were resuspended in 0.1% formic acid to a final concentration of 250 ng/µL.

For quantitative proteomics, 1 µg of trypsin-digested peptides was resuspended in 50 mM TEAB (pH 8.5) and labeled using the TMT 10plex Isobaric Labeling Reagent Set (Thermo Fisher Scientific, 90406). Each sample was labeled with TMT 10plex reagents at a 10:1 label-to-peptide ratio (w/w) for 1 hour at room temperature. The labeling reaction was then quenched by adding 0.2% hydroxylamine. Samples were pooled and then lyophilized. The TMT-labeled peptides were fractionated using the Pierce High pH Reversed-Phase Peptide Fractionation Kit (Thermo Fisher Scientific, 84868) following the manufacturer’s protocol. The fractions were then flash-frozen and lyophilized. Lyophilized peptides were resuspended in 0.2% formic acid to a concentration of 250 ng/μL.

An Orbitrap Exploris mass spectrometer equipped with a FAIMS Pro interface connected to an Easy-nLC 1200 chromatography system (Thermo Fisher Scientific) was used to analyze peptides. An Aurora Ultimate 25 cm column (75 µm × 25 cm, 120 Å) from IonOpticks (Fitzroy, VIC, AUS) was used to separate peptides. A gradient of 0.1% formic acid in water (eluent A) and 0.1% formic acid in 80% acetonitrile (eluent B) was run with the following steps: 1) 5–25% B for 110 min, 2) 25–40% B for 10 min, 3) 40–95% B for 10 min, 4) 95% B over 10 min, 5) 95-2% B for 2 min, 6) 2% B for 2 min, 7) 2-98% B for 2 min, 8) 98% B for 2 min, 9) 98-2% B for 2 min, and 10) 2% B for 2 min, using 0.1% FA. The Orbitrap and FAIMS Pro were operated in positive ion mode with a positive ion voltage of 1800 V, an ion transfer tube temperature of 270°C, and a 4.6 L/min carrier gas flow, using standard FAIMS resolution and compensation voltages of -45 and -65 V. Full scan spectra were acquired in profile mode at a resolution of 60,000 (MS1) and 45,000 (MS2), with a scan range of 400–1400 m/z, custom maximum fill time (50 ms), custom AGC target (300% MS1, 200% MS2), isolation windows of m/z 0.7, intensity threshold of 5.0e^4^, 2–5 charge state, dynamic exclusion of 30 seconds, and 32% HCD collision energy.

Mass spectrometry data were processed with Proteome Discoverer 3.2 (Thermo Fisher Scientific). Sequest HT was used to identify proteins and peptides (Thermo Fisher Scientific). The human protein database (UP000005640) was used as the reference database and the sequence of GFP and each HP1 isoform was added to data processing.

For label-free mass spectrometry, tryptic peptides with up to 2 missed cleavages were allowed. Mass tolerances for precursors were set to 10 ppm and fragments were set to 0.02 Da. The analysis included the following post-translational modifications: dynamic oxidation (+15.995 Da; M), dynamic acetylation (+42.011 Da; N-terminus), dynamic Met-loss (–131.04 Da; M N-terminus), dynamic Met-loss combined with acetylation (–89.03 Da; M N-terminus), and static carbamidomethylation (+57.021 Da; C). For TMT experiments, we included static TMT (+229.163 Da; any N-terminus), and static TMT (+229.163 Da; K). TMT isotope correction values were accounted for (Thermo Fisher; lot no. VK306786). Percolator with an FDR of less than 0.01 was used to filter the identified peptides.

### Identification of peptides from published mass spectrometry data

For peptides identified using Proteome Discoverer 3.2 (Thermo Fisher), the indicated mass spectrometry data were processed as follows. First, Sequest HT was used to identify proteins and peptides (Thermo Fisher Scientific). The human protein database (UP000005640) was used as the reference database and the sequence of each HP1 isoform was added to data processing. For each dataset processing followed what was described in the PRIDE database. Briefly, for PXD014802 GluC was used as the enzyme with up to 3 missed cleavages allowed. Mass tolerances for precursors were set to 10 ppm and fragments were set to 0.02 Da. The analysis included the following post-translational modifications: dynamic oxidation (+15.995 Da; M), dynamic acetylation (+42.011 Da; N-terminus), dynamic methylation (+14.016 Da; K), dynamic dimethyl (+28.031 Da; K), dynamic trimethyl (+42.047 Da; K), dynamic Met-loss (–131.04 Da; M N-terminus), dynamic Met-loss combined with acetylation (–89.03 Da; M N-terminus), and static carbamidomethylation (+57.021 Da; C). For PXD006633 ArgC was used as the enzyme with a maximum of 2 missed cleavages allowed. Mass tolerances for precursors were set to 10 ppm and fragments were set to 0.02 Da. The analysis included the following post-translational modifications: dynamic oxidation (+15.995 Da; M), dynamic dimethyl (+28.031 Da, peptide terminus), dynamic acetylation (+42.011 Da; N-terminus), dynamic Met-loss (–131.04 Da; M N-terminus), dynamic Met-loss combined with acetylation (–89.03 Da; M N-terminus), and static carbamidomethylation (+57.021 Da; C) and dimethyl (+28.031 Da; K). For PXD025053 the mouse protein database (UP000000589), along with the sequences of the mouse HP1 isoforms was used for data processing. Trypsin was used as the enzyme and up to 2 missed cleavages were allowed. Mass tolerances for precursors were set to 10 ppm and fragments were set to 0.02 Da. The analysis included the following post-translational modifications: dynamic oxidation (+15.995 Da; M), dynamic acetylation (+42.011 Da; N-terminus), dynamic methylation (+14.016 Da; K), dynamic dimethyl (+28.031 Da; K), dynamic trimethyl (+42.047 Da; K), dynamic Met-loss (–131.04 Da; M N-terminus), dynamic Met-loss combined with acetylation (–89.03 Da; M N-terminus), and static carbamidomethylation (+57.021 Da; C). For each dataset Percolator with an FDR of less than 0.01 was used to filter the identified peptides.

For dataset PXD010154 and PDC00219, the PepQuery2 webserver was used with the default settings. The data were searched with different tryptic peptides matching to the extension. For PXD058868 peptides were identified with PepQuery Version 2.0.2 with ArgC as the enzyme allowing for up to 2 missed cleavages^41^. Mass tolerances for precursors were set to 20 ppm and fragments were set to 0.02 Da. The analysis included the following post-translational modifications: dynamic oxidation (+15.995 Da; M), dynamic acetylation (+42.011 Da; N-terminus), and static carbamidomethylation (+57.021 Da; C).

### Structure Prediction and Visualization

Structure predictions were carried out on the AlphaFold 3 webserver and structures were visualized with PyMOL 2.0 (Schrodinger LLC).

### Protein Purification

To express and purify all HP1β constructs, *Escherichia coli* BL21-DE3 cells were grown in an incubating shaker at 37 °C and 215 rpm until they reached an OD_600_ of 0.8-1.0. Upon reaching this optical density expression was induced with 0.5 mM Isopropyl β-d-1-thiogalactopyranoside (IPTG) for 14-16 hours at 24 °C and 115 rpm. Cells were then spun down at 4,000 x g (JA-14 Beckman Coulter). Cell pellets were resuspended in lysis buffer (25 mM Tris HCl pH 7.5, 300 mM NaCl, 1 mM BME, 1 mM PMSF) supplemented with a protease inhibitor tablet. Cells were lysed by sonication and the lysate was clarified by centrifugation at 18,000 rpm in a Ti-70 rotor (Beckman Coulter) in an Optima XE-90 ultra-centrifuge (Beckman Coulter). Clarified lysate was passed over a Glutathione-Sepharose column (Glutathione Sepharose 6 fast flow, GE Healthcare) that was equilibrated with wash buffer (25 mM Tris HCl pH 7.5, 300 mM NaCl, 1 mM BME). After the lysate was passed over, the column was washed with 10 column volumes of wash buffer. The proteins were eluted from the column by overnight incubation in lysis buffer containing PreScission protease. Cleaved protein was eluted and then dialyzed into S75 buffer (50 mM Tris HCl pH 8, 150 mM NaCl, 1 mM DTT) at 4 °C for at least 4 hours. After dialysis the protein was passed over a S-75 increase column (Superdex 75 Increase 10/300GL, Cytiva) with S75 buffer as the eluent. Peak fractions were analyzed by Coomassie stained SDS-PAGE gels to determine purity. Pure fractions were then pooled, concentrated, flash frozen and stored at -80°C.

### Isothermal titration calorimetry

Peptides were synthesized by the Swanson Biotechnology Core at Koch Institute for Integrative Cancer Research at MIT. The sequence used for the CAF-1 peptide is N-GILFKGKVPMVVLQDILAVR-C. All proteins were dialyzed into 20 mM Tris pH 7.5 and 150 mM NaCl at 4 °C overnight. Protein concentrations were determined by measuring the absorbance at 280 nm with a spectrophotometer (Nanodrop 2000). Peptides were resuspended to 200 µM in the spent dialysis buffer. Proteins were diluted to 20 µM using the spent dialysis buffer. Measurements were performed using a PEAQ-ITC microcalorimeter (Malvern analytical) at 25 °C with a reference power of 10 µcal/s, and stir speed set to 750 rpm. HP1β constructs were loaded into the chamber and peptides were titrated in over 18 injections. Titration was performed with a 0.4 µL first injection (to remove any air) and all subsequent injections were 2 µL over 4 seconds with 150 second spacing between each injection. The data were analyzed using MicroCal PEAQ-ITC Analysis Software (Malvern), background was corrected using a fitted offset, and data was fit as a single site binding model.

**Supplemental Figure 1.**
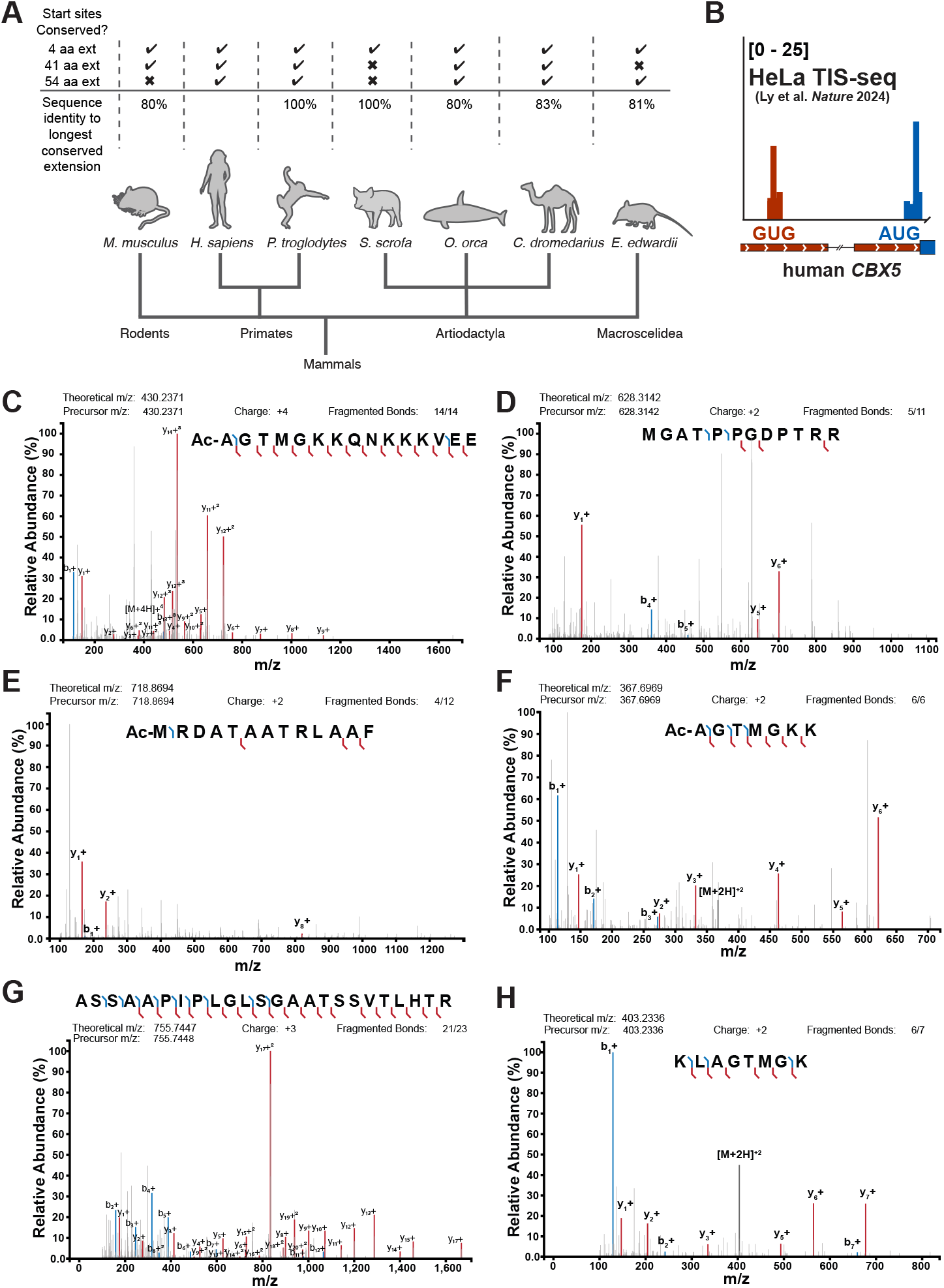
Multiple HP1β isoforms can be produced from human and mouse *CBX1* mRNA. **(A)** A cartoon evolutionary tree highlighting the conservation of each extended start site across mammals. Sequences were identified using tblastn. The sequence identity of the longest conserved extension is compared to the human protein sequence. **(B)** Translation initiation site ribosome profiling read distribution at human *CBX5* alternative start codons. The red peaks are reads at the alternative start sites in the 5’ UTR of the transcript and the blue peaks map to the annotated start site. **(C)-(H)** Annotated MS/MS spectra with the relative abundance plotted against the mass/charge. Each spectrum was generated from raw data using the Interactive Peptide Spectral Annotator^43^. b-ions are shown in blue and y-ions are shown in red. Theoretical monoisotopic m/z values were calculated using Protein Prospector (UCSF).

**Supplemental Figure 2.**
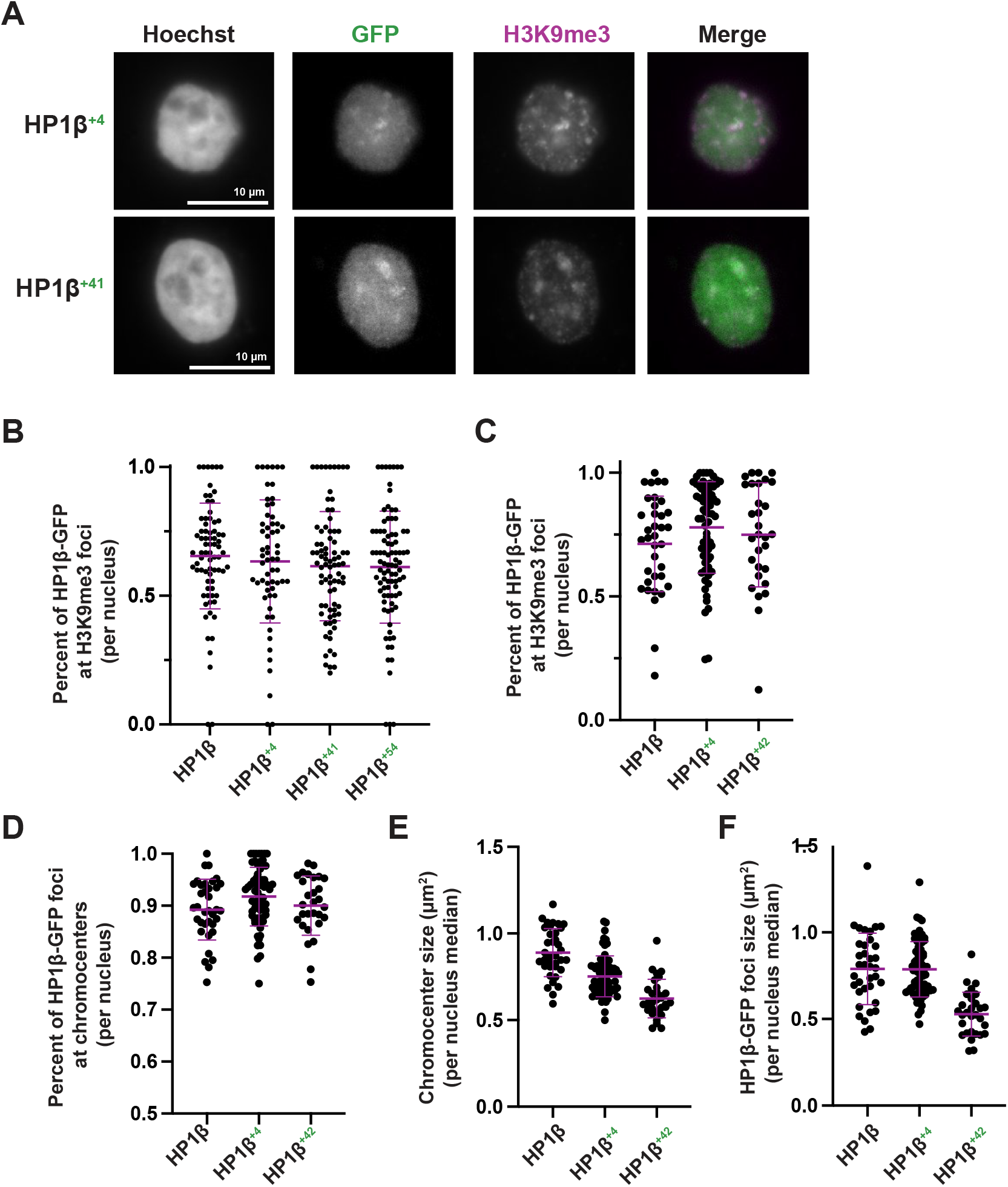
Isoforms of HP1β have differential chromatin localization. **(A)** Representative images of a single 2 µm equatorial slice of HeLa cells ectopically expressing the indicated C-terminally GFP tagged HP1β construct. The Hoechst and H3K9me3 channels are scaled identically, but the GFP channels are scaled individually. **(B)** A dot plot showing the fraction per nucleus of HP1β-GFP foci that overlap with H3K9me3 foci in HeLa cells. Each dot represents a single nucleus. The mean and error bars represent the standard deviation from three biological replicates. **(C)** A dot plot showing the fraction per nucleus of HP1β-GFP foci that overlap with H3K9me3 foci in NIH3T3 cells. Each dot represents a single nucleus. The mean and error bars represent the standard deviation from three biological replicates. **(D)** A dot plot showing the fraction per nucleus of HP1β-GFP foci that overlap with chromocenters in NIH3T3 cells. The mean and error bars represent the standard deviation from three biological replicates. **(E)** A dot plot showing the median size (µm^2^) of chromocenters per nucleus in NIH3T3 cells. Each dot represents a single nucleus. The mean and error bars represent the standard deviation from three biological replicates. **(F)** A dot plot showing the median size (µm^2^) of HP1β-GFP foci per nucleus in NIH3T3 cells. Each dot represents the median size from a single nucleus. The mean and error bars represent the standard deviation from three biological replicates.

**Supplemental Figure 3.**
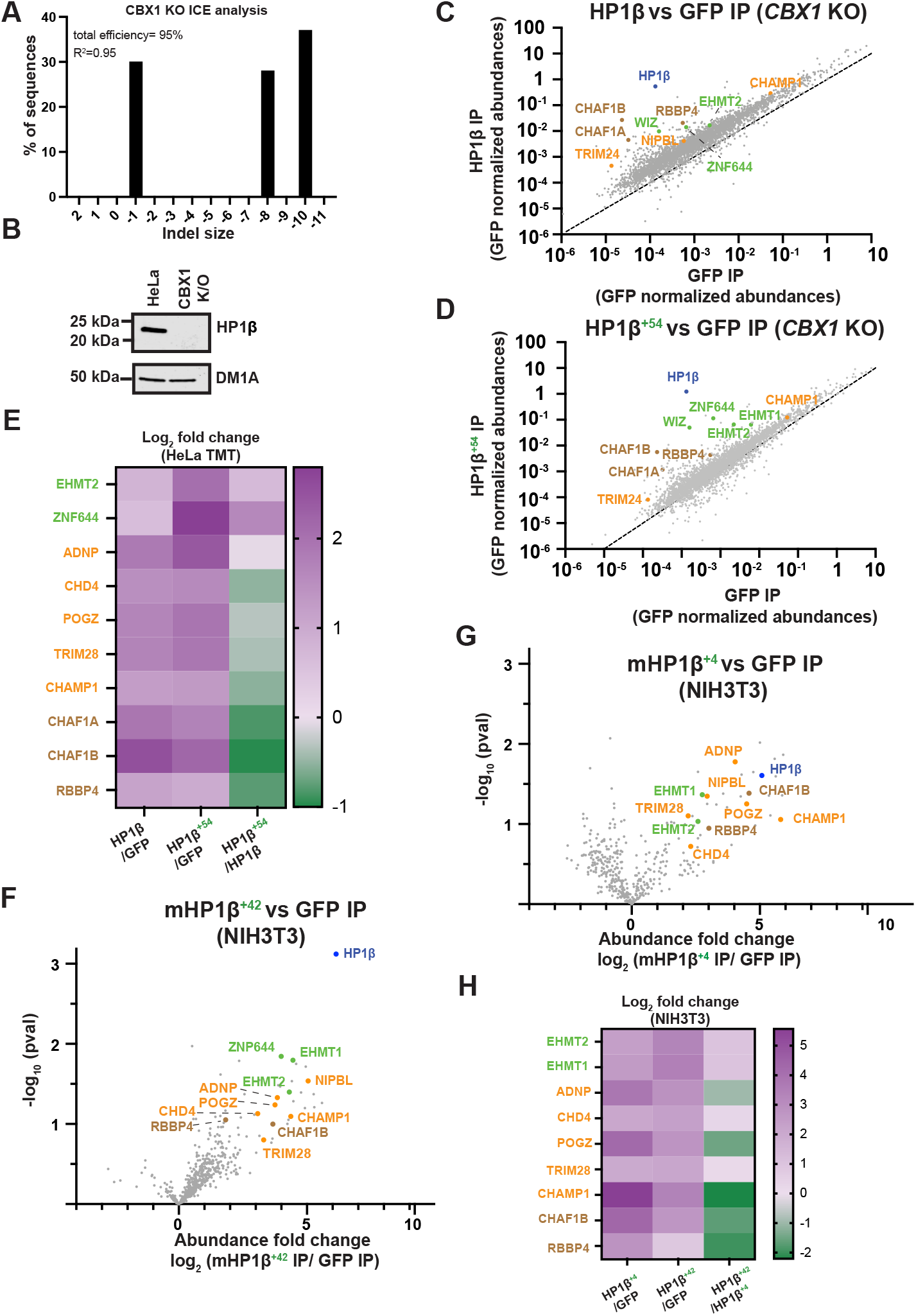
Quantitative mass spectrometry reveals preferential interactions between HP1β isoforms and chromatin effectors. **(A)** Sanger sequencing-based analysis of *CBX1* knockout from a single HeLa clone. ICE (Synthego) was used to deconvolve the Sanger sequencing^44^. **(B)** A western blot showing HP1β expression levels in the indicated cell lines. **(C)-(D)** Scatter plots showing the GFP normalized abundance of the proteins detected in the HP1β^+54^ IP or HP1β IP compared to the GFP IP from HeLa cells in which *CBX1* was targeted using CRISPR/Cas9. HP1β is highlighted in blue, components of the G9a/GLP complex are shown in green, PxVxL containing proteins are shown in orange, and the CAF-1 complex subunits are shown in brown. **(E)** A heatmap showing the log_2_ fold abundance change of selected proteins in the indicated IP experiments from HeLa cells. **(F)-(G)** Volcano plots showing the log_2_ fold change of HP1β^+42^ IP/GFP or HP1β^+4^ IP/GFP IP from TMT-based proteomics experiments in NIH3T3 cells. **(H)** A heatmap showing the log_2_ fold abundance change of selected proteins in the indicated IP experiments.

**Supplemental Figure 4.**
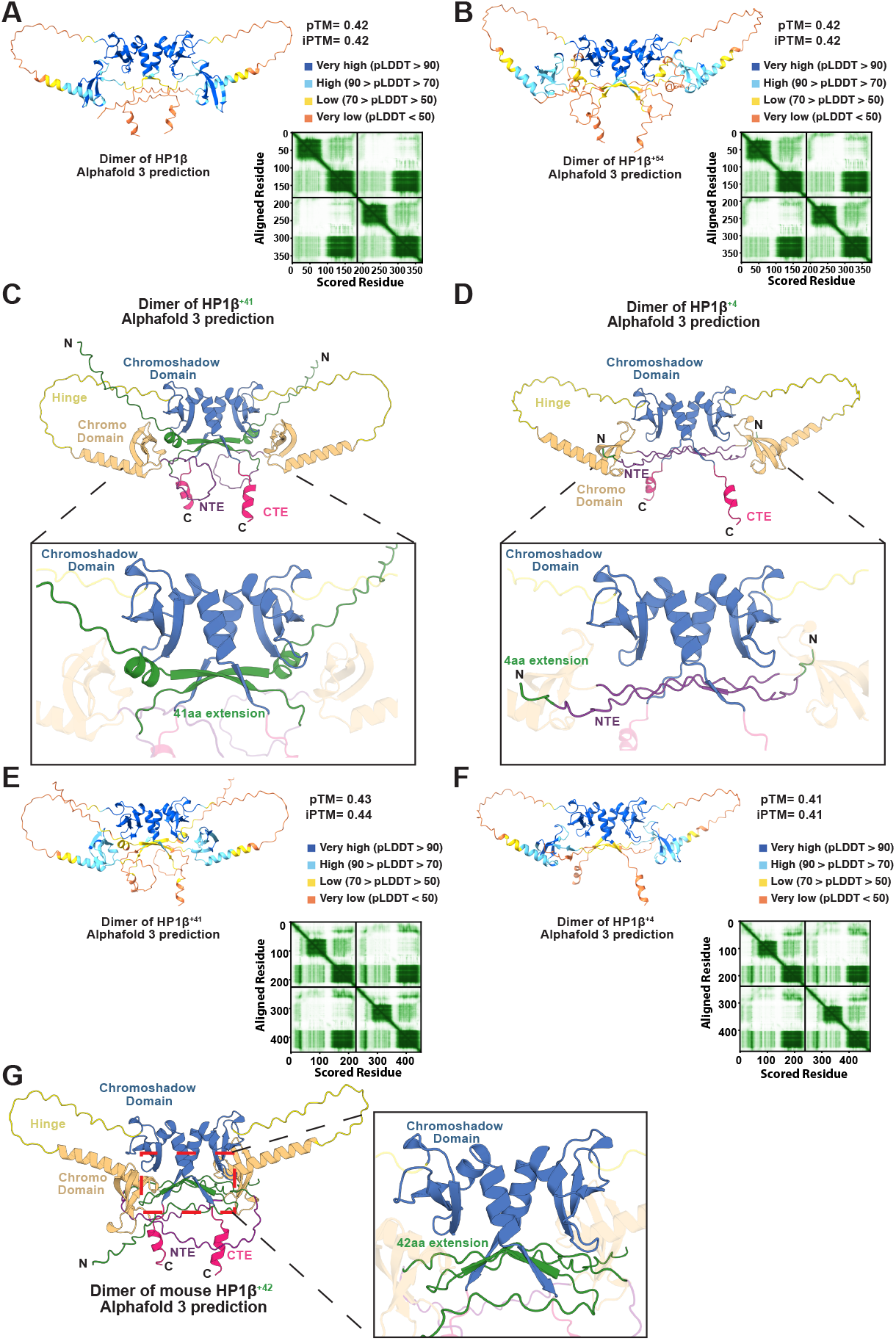
Structural predictions show an intramolecular interaction between the HP1β N-terminal extension and its effector binding groove. **(A)** The predicted structure of the dimer of annotated HP1β colored by the predicted local distance difference test (pLDDT) confidence score generated by AlphaFold 3. The predicted aligned error (PAE) plot generated by AlphaFold 3 is included to the right of the predicted structure. **(B)** The predicted structure of the dimer of HP1β^+54^ colored by the pLDDT confidence score generated by AlphaFold 3. The predicted aligned error (PAE) plot generated by AlphaFold 3 is included to the right of the predicted structure. **(C)** A cartoon representation of the predicted structure of the dimer of HP1β^+41^ that was created using AlphaFold 3. Each chain of the dimer has the same coloring scheme. For each chain the N-terminal extension (NTE) is shown in purple, the chromodomain is shown in orange, the hinge region is shown in yellow, the chromoshadow domain is shown in blue, and the C-terminal extension is shown in pink, and the residues that are appended to the N-terminus by the translational isoform are shown in green. The bottom panel is a zoomed in view of the hydrophobic binding pocket that is formed by the CSD-CSD mediated homodimerization interface. The bottom panel highlights the interaction the extended isoform is predicted to make with the hydrophobic pocket. **(D)** A cartoon representation of the predicted structure of the dimer of HP1β^+4^ that was created using AlphaFold 3. The color scheme is the same as (C). The bottom panel highlights the interaction the extended isoform is predicted to make with the hydrophobic pocket. **(E)** The predicted structure of the dimer of HP1β^+41^ colored by the pLDDT confidence score generated by AlphaFold 3. The PAE plot generated by AlphaFold 3 is included to the right of the predicted structure. **(F)** The predicted structure of the dimer of HP1β^+4^ colored by the pLDDT confidence score generated by AlphaFold 3. The PAE plot generated by AlphaFold 3 is included to the right of the predicted structure. **(G)** A cartoon representation of the predicted structure of the dimer of mouse HP1β^+42^ that was created using AlphaFold 3. The color scheme is the same as (C). The panel to the right highlights the interaction the extended isoform is predicted to make with the hydrophobic pocket.

**Supplemental Figure 5.**
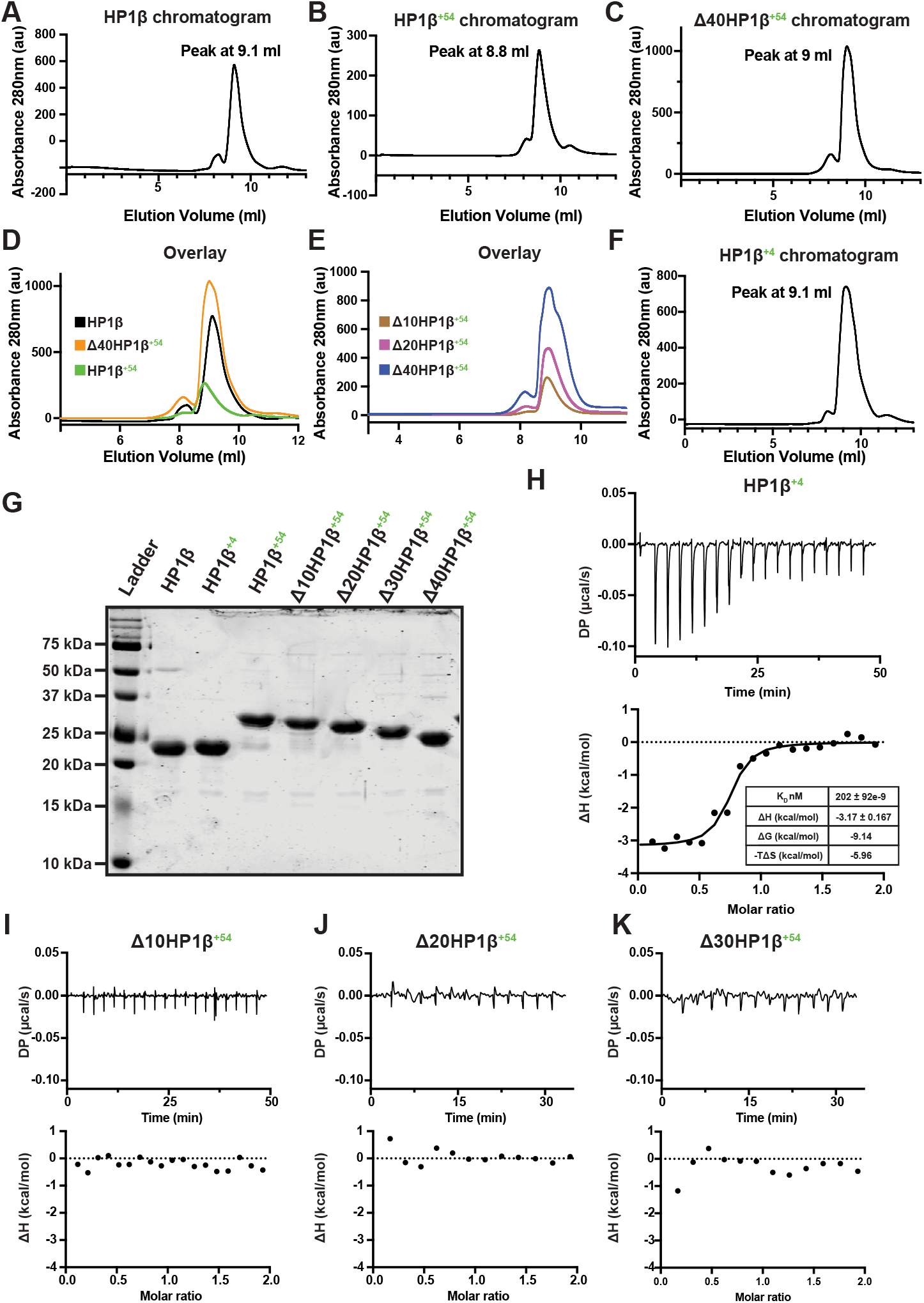
N-terminally extended isoforms of HP1β fail to bind to PxVxL containing peptide. **(A)-(F)** Representative chromatograms or overlaid chromatograms of the indicated HP1β construct. **(G)** A Coomassie stained SDS-PAGE gel of the pooled and concentrated fraction of each HP1β construct used for ITC. **(H)-(K)** Representative raw thermograms (top) and integrated heats fit to a single-site model (bottom). For each panel a CAF-1 peptide^4^ was titrated into the cell containing the indicated HP1β construct.

**Supplemental Table 1. Mass Spectrometry Data.**

## References

1. Jacobs, S. A. & Khorasanizadeh, S. Structure of HP1 chromodomain bound to a lysine 9-methylated histone H3 tail. Science 295, 2080–2083 (2002). 10.1126/science.1069473

2. Bannister, A. J. et al. Selective recognition of methylated lysine 9 on histone H3 by the HP1 chromo domain. Nature 410, 120–124 (2001). 10.1038/35065138

3. Lachner, M., O’Carroll, D., Rea, S., Mechtler, K. & Jenuwein, T. Methylation of histone H3 lysine 9 creates a binding site for HP1 proteins. Nature 410, 116–120 (2001). 10.1038/35065132

4. Brasher, S. V. et al. The structure of mouse HP1 suggests a unique mode of single peptide recognition by the shadow chromo domain dimer. EMBO J 19, 1587–1597 (2000). 10.1093/emboj/19.7.1587

5. Thiru, A. et al. Structural basis of HP1/PXVXL motif peptide interactions and HP1 localisation to heterochromatin. EMBO J 23, 489–499 (2004). 10.1038/sj.emboj.7600088

6. Colmenares, S. U. et al. Expanding the HP1a-binding consensus and molecular grammar for heterochromatin assembly. Nucleic Acids Res 53 (2025). 10.1093/nar/gkaf976

7. Fischle, W. et al. Molecular basis for the discrimination of repressive methyl-lysine marks in histone H3 by Polycomb and HP1 chromodomains. Genes Dev 17, 1870–1881 (2003). 10.1101/gad.1110503

8. Canzio, D., Larson, A. & Narlikar, G. J. Mechanisms of functional promiscuity by HP1 proteins. Trends Cell Biol 24, 377–386 (2014). 10.1016/j.tcb.2014.01.002

9. Pak, D. T. et al. Association of the origin recognition complex with heterochromatin and HP1 in higher eukaryotes. Cell 91, 311–323 (1997). 10.1016/s0092-8674(00)80415-8

10. Quivy, J. P., Gerard, A., Cook, A. J., Roche, D. & Almouzni, G. The HP1-p150/CAF-1 interaction is required for pericentric heterochromatin replication and S-phase progression in mouse cells. Nat Struct Mol Biol 15, 972–979 (2008). 10.1038/nsmb.1470

11. Vakoc, C. R., Mandat, S. A., Olenchock, B. A. & Blobel, G. A. Histone H3 lysine 9 methylation and HP1gamma are associated with transcription elongation through mammalian chromatin. Mol Cell 19, 381–391 (2005). 10.1016/j.molcel.2005.06.011

12. Schoelz, J. M. & Riddle, N. C. Functions of HP1 proteins in transcriptional regulation. Epigenetics Chromatin 15, 14 (2022). 10.1186/s13072-022-00453-8

13. Nielsen, S. J. et al. Rb targets histone H3 methylation and HP1 to promoters. Nature 412, 561–565 (2001). 10.1038/35087620

14. Ainsztein, A. M., Kandels-Lewis, S. E., Mackay, A. M. & Earnshaw, W. C. INCENP centromere and spindle targeting: identification of essential conserved motifs and involvement of heterochromatin protein HP1. J Cell Biol 143, 1763–1774 (1998). 10.1083/jcb.143.7.1763

15. Hiragami-Hamada, K. et al. Dynamic and flexible H3K9me3 bridging via HP1beta dimerization establishes a plastic state of condensed chromatin. Nat Commun 7, 11310 (2016). 10.1038/ncomms11310

16. Smothers, J. F. & Henikoff, S. The HP1 chromo shadow domain binds a consensus peptide pentamer. Curr Biol 10, 27–30 (2000). 10.1016/s0960-9822(99)00260-2

17. Bosch-Presegue, L. et al. Mammalian HP1 Isoforms Have Specific Roles in Heterochromatin Structure and Organization. Cell Rep 21, 2048–2057 (2017). 10.1016/j.celrep.2017.10.092

18. Aucott, R. et al. HP1-beta is required for development of the cerebral neocortex and neuromuscular junctions. J Cell Biol 183, 597–606 (2008). 10.1083/jcb.200804041

19. Brown, J. P. et al. Correction: HP1gamma function is required for male germ cell survival and spermatogenesis. Epigenetics Chromatin 5, 18 (2012). 10.1186/1756-8935-5-18

20. Ayoub, N., Jeyasekharan, A. D., Bernal, J. A. & Venkitaraman, A. R. HP1-beta mobilization promotes chromatin changes that initiate the DNA damage response. Nature 453, 682–686 (2008). 10.1038/nature06875

21. Maison, C. et al. SUMOylation promotes de novo targeting of HP1alpha to pericentric heterochromatin. Nat Genet 43, 220–227 (2011). 10.1038/ng.765

22. Lee, S. et al. Global mapping of translation initiation sites in mammalian cells at single-nucleotide resolution. Proc Natl Acad Sci U S A 109, E2424–2432 (2012). 10.1073/pnas.1207846109

23. Ingolia, N. T., Lareau, L. F. & Weissman, J. S. Ribosome profiling of mouse embryonic stem cells reveals the complexity and dynamics of mammalian proteomes. Cell 147, 789–802 (2011). 10.1016/j.cell.2011.10.002

24. Ly, J. et al. Alternative start codon selection shapes mitochondrial function and rare human diseases. Mol Cell 85, 4198–4214 e4111 (2025). 10.1016/j.molcel.2025.10.013

25. Tsang, M. J. & Cheeseman, I. M. Alternative CDC20 translational isoforms tune mitotic arrest duration. Nature 617, 154–161 (2023). 10.1038/s41586-023-05943-7

26. Marescal, O. & Cheeseman, I. M. A machine learning model predicts protein stability of annotated and alternate protein isoforms. bioRxiv (2026). 10.64898/2026.07.16.739020

27. Ly, J. et al. Nuclear release of eIF1 restricts start-codon selection during mitosis. Nature 635, 490–498 (2024). 10.1038/s41586-024-08088-3

28. Legartova, S. et al. DNA Damage Changes Distribution Pattern and Levels of HP1 Protein Isoforms in the Nucleolus and Increases Phosphorylation of HP1beta-Ser88. Cells 8 (2019). 10.3390/cells8091097

29. Na, C. H. et al. Discovery of noncanonical translation initiation sites through mass spectrometric analysis of protein N termini. Genome Res 28, 25–36 (2018). 10.1101/gr.226050.117

30. Qin, W. et al. Phosphorylation of the HP1beta hinge region sequesters KAP1 in heterochromatin and promotes the exit from naive pluripotency. Nucleic Acids Res 49, 7406–7423 (2021). 10.1093/nar/gkab548

31. Ju, S. et al. Implementing N-terminomics and machine learning to probe Nt-arginylation. Nat Commun 17, 172 (2025). 10.1038/s41467-025-66883-6

32. Saksouk, N., Simboeck, E. & Dejardin, J. Constitutive heterochromatin formation and transcription in mammals. Epigenetics Chromatin 8, 3 (2015). 10.1186/1756-8935-8-3

33. Maison, C., Quivy, J. P., Probst, A. V. & Almouzni, G. Heterochromatin at mouse pericentromeres: a model for de novo heterochromatin formation and duplication during replication. Cold Spring Harb Symp Quant Biol 75, 155–165 (2010). 10.1101/sqb.2010.75.013

34. Burton, A. et al. Heterochromatin establishment during early mammalian development is regulated by pericentromeric RNA and characterized by non-repressive H3K9me3. Nat Cell Biol 22, 767–778 (2020). 10.1038/s41556-020-0536-6

35. Jagannathan, M., Cummings, R. & Yamashita, Y. M. The modular mechanism of chromocenter formation in Drosophila. Elife 8 (2019). 10.7554/eLife.43938

36. Sokolova, V. et al. Structural mechanism of HP1α-dependent transcriptional repression and chromatin compaction. Structure 32, 2094–2106 e2096 (2024). 10.1016/j.str.2024.09.013

37. Richart, A. N., Brunner, C. I., Stott, K., Murzina, N. V. & Thomas, J. O. Characterization of chromoshadow domain-mediated binding of heterochromatin protein 1alpha (HP1alpha) to histone H3. J Biol Chem 287, 18730–18737 (2012). 10.1074/jbc.M111.337204

38. Sales-Gil, R. & Vagnarelli, P. How HP1 Post-Translational Modifications Regulate Heterochromatin Formation and Maintenance. Cells 9 (2020). 10.3390/cells9061460

39. Canzio, D. et al. A conformational switch in HP1 releases auto-inhibition to drive heterochromatin assembly. Nature 496, 377–381 (2013). 10.1038/nature12032

40. Cheeseman, I. M. & Desai, A. A combined approach for the localization and tandem affinity purification of protein complexes from metazoans. Sci STKE 2005, p1 (2005). 10.1126/stke.2662005pl1

41. Wen, B. & Zhang, B. PepQuery2 democratizes public MS proteomics data for rapid peptide searching. Nat Commun 14, 2213 (2023). 10.1038/s41467-023-37462-4

42. Abramson, J. et al. Accurate structure prediction of biomolecular interactions with AlphaFold 3. Nature 630, 493–500 (2024). 10.1038/s41586-024-07487-w

43. Brademan, D. R., Riley, N. M., Kwiecien, N. W. & Coon, J. J. Interactive Peptide Spectral Annotator: A Versatile Web-based Tool for Proteomic Applications. Mol Cell Proteomics 18, S193–S201 (2019). 10.1074/mcp.TIR118.001209

44. Conant, D. et al. Inference of CRISPR Edits from Sanger Trace Data. CRISPR J 5, 123–130 (2022). 10.1089/crispr.2021.0113

